# MipSScs: Artificial neural network-based data integration of 2D/3D single-cell spatial RNA sequence data from virus-infected human cerebral organoids

**DOI:** 10.64898/2026.06.08.727881

**Authors:** Anusha D Doddi, Pepper Dawes, Yingleong Chan, Elaine T. Lim

## Abstract

There is interest in the use of recent single-cell spatial transcriptomic technologies to gain biological insights into disease mechanisms. Previously, we characterized the use of herpes simplex virus 1 (HSV-1) induced neuroinflammation in 2D dissociated cells from human cerebral organoids (dcOrgs) to model molecular and transcriptomic readouts associated with Alzheimer’s disease (AD). In this work, we generated two datasets by using single-cell non-spatial RNA sequencing and single-cell spatial RNA sequencing technologies on HSV-1-infected 2D dcOrgs and HSV-1-infected 3D cerebral organoids (cOrgs). We conducted cell type assignment for the cells in the 2D dcOrgs and 3D cOrgs, by using single-cell non-spatial RNA sequence data from human fetal brains and adult post-mortem brains, to infer the transcriptomic effects of AD-associated *in-vitro* perturbations through viral infections linked to AD.

We evaluated computational and machine learning methods, including the use of multi-layer perceptrons (MLPs), and we used cross-2D/3D platform comparisons as a benchmark to evaluate the artificial neural network models. In the process, we found that the use of MLPs can lead to high validation rates for assigning cell type identities from 2D and 3D human cerebral organoids to cell types found in human adult post-mortem brain samples. Furthermore, the use of these technologies and systems enabled the identification of pseudotime trajectories and cell clusters associated with the viral transcriptional life cycle. We identified several cell types, including endothelial cells and astrocytes, with significantly more clustered cell-cell nearest neighbor distances in infected 3D cOrgs compared to mock 3D cOrgs. Permutation tests revealed that these differences in nearest neighbor distances are unlikely to be driven by overall structural differences between individual infected 3D cOrgs and mock 3D cOrgs, such as differences in the density of cells.

Given that there are more large-scale single-cell non-spatial (2D) RNA sequence datasets that had been generated from human post-mortem brain samples, compared to single-cell spatial (3D) RNA sequence datasets from human post-mortem brain samples, the development of data integration approaches by using artificial neural networks such as MLPs, across 2D and 3D single-cell transcriptomics datasets generated from human post-mortem brain samples and human *in-vitro* systems such as brain organoids is likely to be critical to gain novel insights into neurodegenerative diseases such as AD.

## INTRODUCTION

There is interest in using single-cell spatial RNA sequencing and single-cell non-spatial RNA technologies to identify disease-associated signatures in Alzheimer’s disease (AD) and other neurodegenerative diseases^1–7^. We used the herpes simplex virus 1 (HSV-1) to induce neuroinflammation in 2D dissociated cells from human cerebral organoids (dcOrgs) that recapitulated molecular features associated with AD^8^. These AD-associated molecular features were accompanied by transcriptomics signatures identified from bulk RNA sequence data and single-cell RNA sequence data (scRNA-seq) from the HSV-1-infected 2D dcOrgs, that could be used to identify key target genes associated with AD. In parallel, we generated a new single-cell spatial RNA sequence data from HSV-1-infected 3D cerebral organoids (that we abbreviated as cOrgs). These datasets enable us to evaluate the use of data integration from 2D dcOrgs and 3D cOrgs with a viral perturbation that led to AD-associated cell type and molecular changes that had been previously reported^8–14^.

Unsupervised machine learning (ML)-based approaches, semi-supervised ML-based approaches and generative artificial intelligence (AI) based approaches to map single-cell data to reference atlases had shown great success^15–20^. Here, we used a type of artificial neural network, multi-layer perceptron (MLP) model, to conduct supervised learning to address a key challenge in cell type assignment from human cerebral organoid data to cell types found in human fetuses, brain tissue samples from children or post-mortem brain tissues from adults, and across 2D to 3D single-cell spatial and non-spatial technologies. We found that the use of MLP models to assign cell types in human cerebral organoids by using single-cell RNA sequence (scRNA-seq) data from adult post-mortem brain samples, led to high cross-2D/3D platform validation rates and cross-reference validation rates.

We further conducted data integration and comparative analyses of single cell transcriptome-wide data from the 2D and 3D systems and technologies to identify key similarities and differences for a comprehensive interrogation of neuroinflammation-associated transcriptomics signatures and found that there were differences in cell type tropism for viral transcript abundance in the 2D dcOrgs and 3D cOrgs systems – a question that cannot be easily addressed without the use of these technologies and computational methods development. Furthermore, we found that there were significant differences in cell-cell nearest neighbor distances for cell types such as endothelial cells and astrocytes in infected 3D cOrgs versus mock 3D cOrgs. We term our suite of tools as MipSScs for “Microphysiological Systems Single cell spatial”.

## RESULTS

### Number of transcripts and percentage of infected cells in HSV-1-infected 2D dcOrgs and 3D cOrgs

We differentiated 5-month-old 2D dcOrgs from a single donor-derived induced pluripotent stem cell (iPSC) by using prior protocols^8,21,22^. We prepared 3 independent samples of mock-infected dcOrgs and another 3 independent samples of HSV-1-infected dcOrgs (an initial 1 million cells in each sample), fixed the cells with 4% paraformaldehyde (PFA) and counted 20,000 single cells from each of the 6 samples, followed by library preparation using the Parse Evercode v2 kit. In parallel, we conducted HSV-1 infection in 3-month-old 3D cOrgs, fixed the cOrgs with 4% PFA and embedded 6 individual cOrgs in an Optimal Cutting Temperature (OCT) cryoblock. In total, we generated 3 independent cryoblocks with 18 mock cOrgs and 3 independent cryoblocks with 18 HSV-1-infected cOrgs, resulting in a total of 36 cOrgs. We obtained a 10μm section from each of the 6 cryoblocks for sample and library preparation with the Stereo-seq v1 kit, leading to 6 independent samples from these 36 cOrgs (Figure 1).

**Figure 1:**
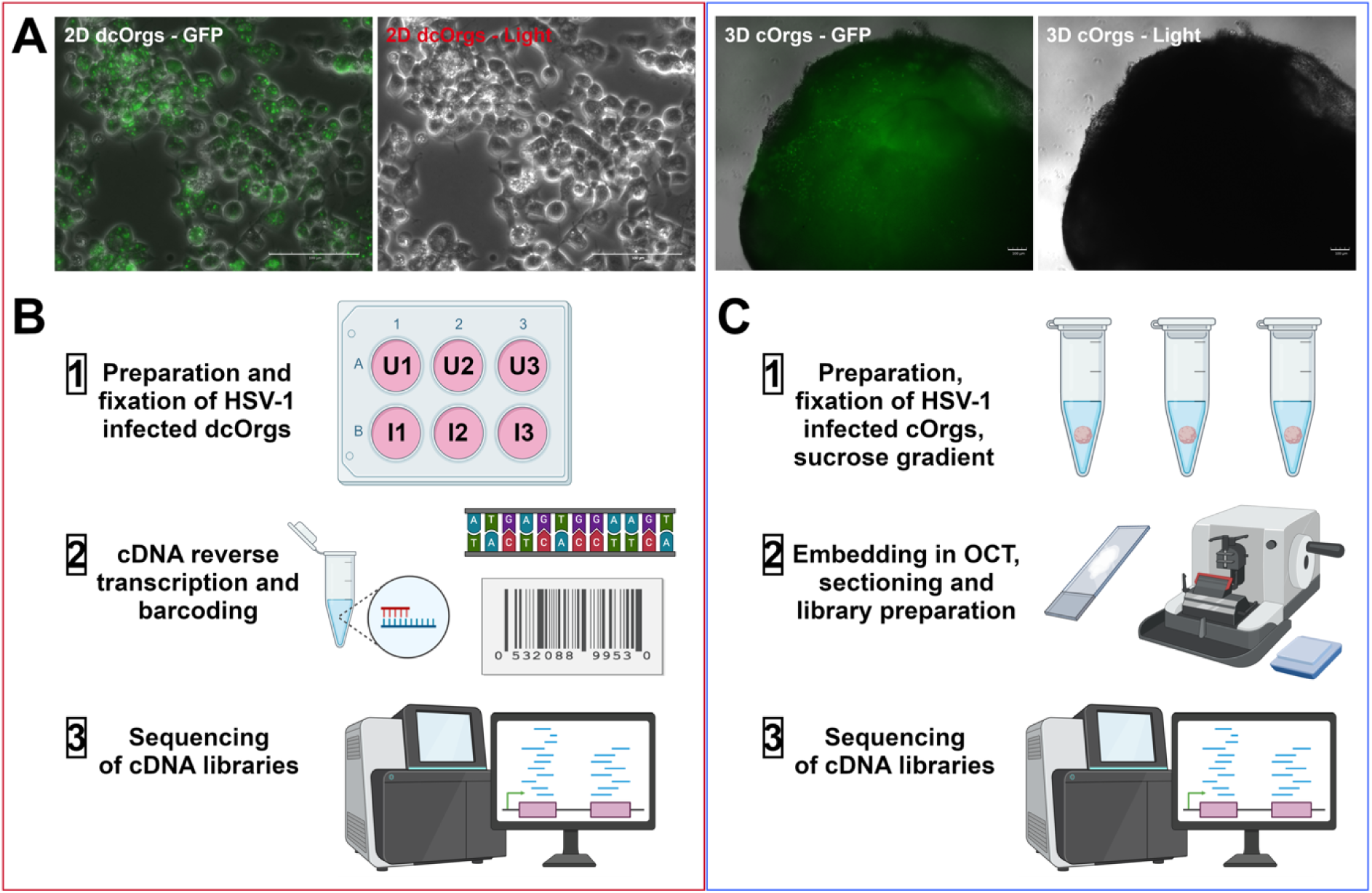
Schematic of the single-cell RNA-seq preparation on HSV-1-infected versus mock 2D dcOrgs and single-cell spatial RNA-seq preparation on HSV-1-infected versus mock 3D cOrgs. (A) Microscope images of HSV-1-infected 2D dcOrgs and HSV-1-infected 3D cOrgs. (B) Sample and library preparation for single-cell RNA sequencing on HSV-1-infected versus mock 2D dcOrgs. (C) Sample and library preparation for single-cell spatial RNA sequencing on HSV-1-infected versus mock 3D cOrgs.

After quality control and data filtering, we obtained 2,815 to 3,800 single cells with 17,102 to 18,524 unique protein-coding transcripts across the 6 sets of 2D dcOrgs and 81-87% of the cells were identified to contain viral transcripts using our thresholds (Extended Data Figures 1-2, Table 1). From each of the 6 sets of 3D cOrg slices, we obtained 6,152 to 25,718 single cells with 17,968 to 18,849 unique protein-coding transcripts and 75-89% of the cells were identified to contain viral transcripts using our thresholds (Table 1).

**Table 1:** Summary of our samples with data generation using two single-cell technologies.

| 2D or 3D | Sample | Infection status | Platform | Total num of cells | Num and % of cells with viral transcripts | Median nFeature SCT | Total nCount RNA |
| --- | --- | --- | --- | --- | --- | --- | --- |
| 2D dcOrg | Mock_dcOrgsP1 | Mock | Evercode | 3,031 | 0 | 11,916 | 40,166,949 |
|  | Mock_dcOrgsP2 | Mock | Evercode | 2,815 | 0 | 10,993 | 35,660,877 |
|  | Mock_dcOrgsP3 | Mock | Evercode | 3,125 | 0 | 12,886 | 44,697,513 |
|  | Inf_dcOrgsP1 | Infected | Evercode | 3,800 | 3,061 (81%) | 2,713 | 16,593,725 |
|  | Inf_dcOrgsP2 | Infected | Evercode | 3,366 | 2,899 (86%) | 3,055 | 15,768,022 |
|  | Inf_dcOrgsP3 | Infected | Evercode | 2,905 | 2,519 (87%) | 3,547 | 15,840,599 |
| **3D**<br>**cOrg** | Mock_cOrgsS1 | Mock | Stereo-seq | 8,515 | 0 | 1,920 | 19,315,060 |
|  | Mock_cOrgsS2 | Mock | Stereo-seq | 17,665 | 0 | 2,022 | 41,694,166 |
|  | Mock_cOrgsS3 | Mock | Stereo-seq | 25,718 | 0 | 1,996 | 56,202,051 |
|  | Inf_cOrgsS1 | Infected | Stereo-seq | 6,152 | 4,736 (77%) | 2,437 | 18,189,951 |
|  | Inf_cOrgsS2 | Infected | Stereo-seq | 12,077 | 9,090 (75%) | 1,847 | 27,524,489 |
|  | Inf_cOrgsS3 | Infected | Stereo-seq | 12,315 | 11,054 (89%) | 2,336 | 34,569,481 |

Details of our samples (mock or HSV-1-infected, 2D dcOrgs or 3D cOrgs), with the number of cells (total number of cells and number of cells with viral transcripts), number of transcripts and number of reads for each sample.

### A consensus approach assigned 36% of cells to a fetal cell type and 7% of cells to an adult cell type

We mapped the identities of individual cells from our 2D and 3D scRNA-seq data to two published scRNA-seq reference datasets generated from fetal brains and brain organoids (we termed this dataset as fetal reference)^23^ and adult post-mortem brains by the Allen Institute (we termed this dataset as adult reference)^24^, by using three cell type assignment tools (Seurat^25,26^, SingleR^27^, and scPred^28^). We used a “consensus” approach through majority voting of the cell type assignment from the three tools. By using the consensus approach on the 2D data (19,042 cells in total), we found that there were 6,873 cells (36.1% of all cells) that had a cell type assignment with the fetal reference and 1,392 cells (7.3% of all cells) that had a cell type assignment with the adult reference (Table 2). By using the consensus approach on the 3D data (82,442 cells in total), we found that there were 20,084 cells (24.4% of all cells) that had a cell type assignment to the fetal reference and 9,491 cells (11.5% of all cells) that had a cell type assignment to the adult reference. These results indicated that in both the 2D and 3D data, there were discrepancies in cell type assignment across the three tools for most of the cells when conducting cell type assignment to the adult reference.

**Table 2:**
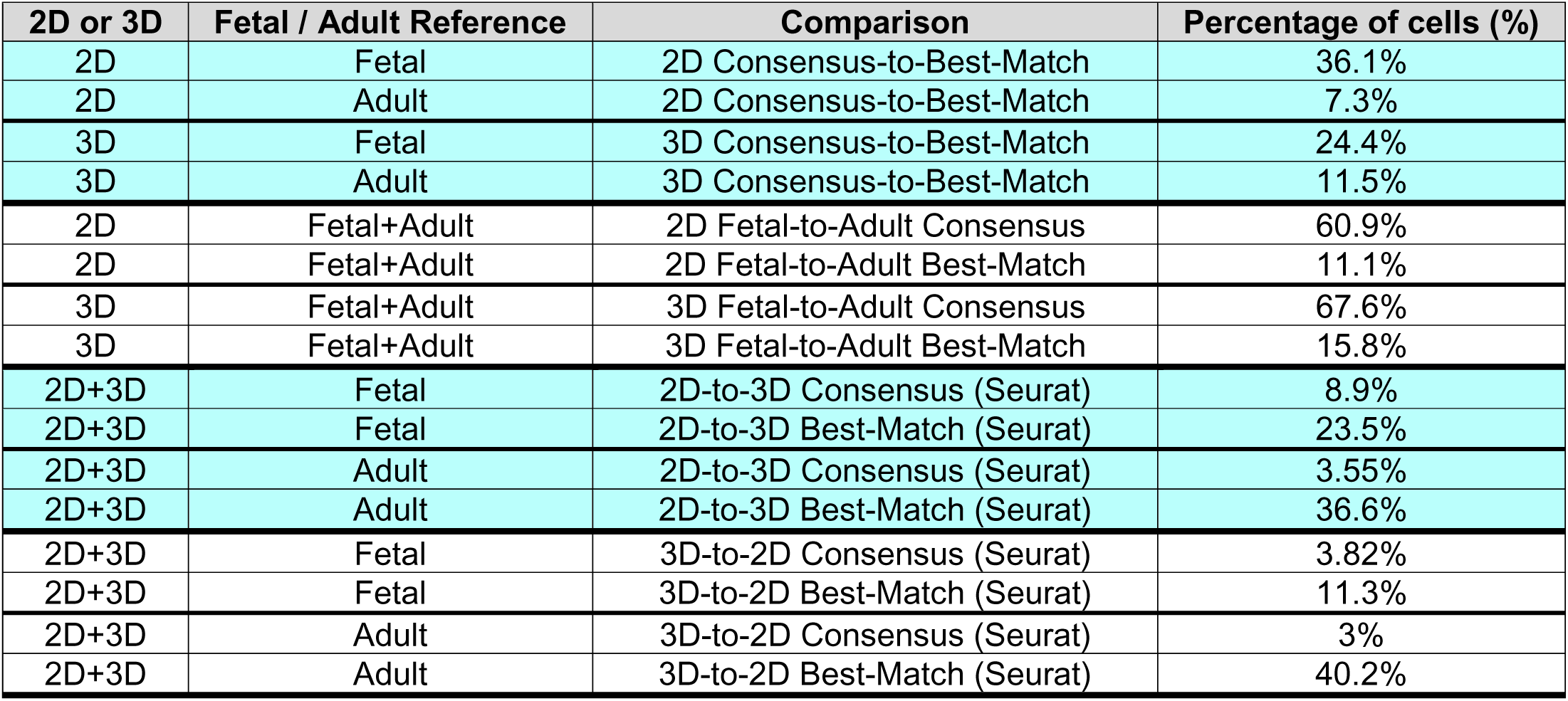
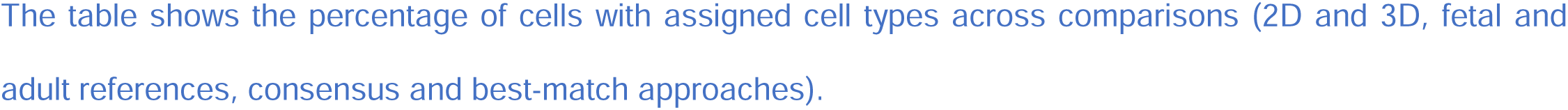
Percentage of cells with assigned cell types across multiple comparisons.

| 2D or 3D | Fetal / Adult Reference | Comparison | Percentage of cells (%) |
| --- | --- | --- | --- |
| 2D | Fetal | 2D Consensus-to-Best-Match | 36.1% |
| 2D | Adult | 2D Consensus-to-Best-Match | 7.3% |
| 3D | Fetal | 3D Consensus-to-Best-Match | 24.4% |
| 3D | Adult | 3D Consensus-to-Best-Match | 11.5% |
| 2D | Fetal+Adult | 2D Fetal-to-Adult Consensus | 60.9% |
| 2D | Fetal+Adult | 2D Fetal-to-Adult Best-Match | 11.1% |
| 3D | Fetal+Adult | 3D Fetal-to-Adult Consensus | 67.6% |
| 3D | Fetal+Adult | 3D Fetal-to-Adult Best-Match | 15.8% |
| 2D+3D | Fetal | 2D-to-3D Consensus (Seurat) | 8.9% |
| 2D+3D | Fetal | 2D-to-3D Best-Match (Seurat) | 23.5% |
| 2D+3D | Adult | 2D-to-3D Consensus (Seurat) | 3.55% |
| 2D+3D | Adult | 2D-to-3D Best-Match (Seurat) | 36.6% |
| 2D+3D | Fetal | 3D-to-2D Consensus (Seurat) | 3.82% |
| 2D+3D | Fetal | 3D-to-2D Best-Match (Seurat) | 11.3% |
| 2D+3D | Adult | 3D-to-2D Consensus (Seurat) | 3% |
| 2D+3D | Adult | 3D-to-2D Best-Match (Seurat) | 40.2% |

The table shows the percentage of cells with assigned cell types across comparisons (2D and 3D, fetal and adult references, consensus and best-match approaches).

Next, we used a “best-match” approach by taking the unassigned cells where the cell types could not be mapped with the consensus approach and designated cell types with the highest confidence scores from the three tools to these unassigned cells. With the best-match approach, all cells in the 2D and 3D data had a cell type assignment. We compared the cells that were assigned cell types by the consensus approach on the 2D and 3D data across the fetal and adult references and found that most cells were assigned equivalent cell types across the fetal and adult references (2D: 60.9% and 3D: 67.6%, Table 2). Conversely, low percentages of cells were assigned equivalent cell types across the fetal and adult references by using the best-match approach (2D: 11.1% and 3D: 15.8%), as shown in Table 2.

The overall percentages of cells that were calculated may be driven by large variances in the percentages calculated for individual cell types. To evaluate this possibility, we evaluated the per-cell type assignment for each of the 13 cell types in the fetal reference and each of the 26 cell types in the adult reference. We found that the individual cell type percentages for 2D consensus-to-best-match ranged from 2% to 63.4% mapped to the fetal reference and from 1.6% to 13.8% mapped to the adult reference (Extended Data Figures 3A-B) and the individual cell type accuracies for 3D consensus-to-best-match ranged from 1.5% to 45.8% mapped to the fetal reference and from 0.36% to 35.7% mapped to the adult reference (Extended Data Figures 3C-D). These results indicate that there were large variances in the percentages of cells for individual cell types.

### Seurat label transfer with best-match assignment from 2D-to-3D data had 23.5% and 36.6% accuracies mapped to fetal and adult references respectively

To evaluate the use of one of the three cell type assignment tools (Seurat) for label transfer of cell types across the 2D and 3D platforms, we used the consensus or best-match assigned cell types in the 2D data to re-assign cell types in the 3D data (and from the 3D data to the 2D data), and evaluated the percentage of these re-assigned cell types that were the same as the best-match assigned cell types. We found that there were lower percentages of cells that were mapped to the same cell types across 2D/3D platforms by conducting label transfer with the consensus assigned cell types (2D-to-3D fetal = 8.9%, 2D-to-3D adult = 3.55%, 3D-to-2D fetal = 3.82%, 3D-to-2D adult = 3%, Table 2), compared to label transfer with the best-match assigned cell types (2D-to-3D fetal = 23.5%, 2D-to-3D adult = 36.6%, 3D-to-2D fetal = 11.3%, 3D-to-2D adult = 40.2%, Table 2). These results indicate that the best-match approach, compared to the use of the consensus approach, led to higher accuracies for cross-platform cell type assignment across the 2D/3D platforms for both the fetal and adult references, possibly because of the larger numbers of cells with assigned cell types by using the best-match approach compared to the numbers of cells with assigned cell types by using the consensus approach.

We evaluated the per-cell type accuracies of using Seurat for label transfer to map each cell across the 2D/3D platforms with the best-match assigned cell types and found that for specific cell types, there were higher percentages of cells that were assigned to the same cell types with the fetal reference (maximum: 50.6% for cortical neurons, Extended Data Figure 4A), when compared to mapping with the adult reference (maximum: 18.2% for vascular leptomeningeal cells or VLMCs, Extended Data Figure 4B). These results are expected, given that the single-cell transcriptomic profiles in the cerebral organoids are more similar to transcriptomic profiles found in human fetal brains than human childhood or adult brains^29^.

### Evaluation of MLP models shows higher overall accuracies and per-cell type accuracies compared to the consensus cell type assignment approach

There are advantages of using MLPs to develop predictive classifiers, such as the ability to learn complex non-linear patterns. We sought to evaluate whether the use of MLPs to conduct cell type assignment on the 2D and 3D data mapped to the fetal and adult references, can lead to improved consistencies in cell type assignment. The 2D and 3D data was split into 9 independent sets of single cells for each dataset (true uninfected or TU cells from the mock samples, true infected or TI cells from the HSV-1-infected samples and pseudo-uninfected or PU cells from the HSV-1-infected samples)^8^. To evaluate the accuracy of the MLP models, we used a bootstrapping approach to leave one of the 9 sets out from training the MLP model and calculated the accuracies of cell type assignment by using the set that left out from training. We evaluated several MLP models with 1 to 3 layers and with 1 to 2,000 nodes. We found that the maximum accuracy for the 2D fetal best-match assignment was 49% and the maximum accuracy for the 2D adult best-match assignment was 41% (Extended Data Figure 5). These accuracies were higher than the consensus assignment compared to best-match assignment on the 2D fetal and adult references (36.1% and 7.3% respectively, Table 2).

The high overall accuracies that we had found by using the MLP models may be driven by specific cell types with more conserved transcriptomic profiles across references or platforms or may be driven by cell types that were more abundant in our datasets. We further evaluated the per-cell type training accuracies of the MLP models and found that the per-cell type training accuracies for the 2D fetal best-match assignment were 72-75% in the 1-layer, 2-layer and 3-layer models with 1,000 nodes for neuroepithelial cells (Extended Data Figures 6A-C) and the per-cell type training accuracies for the 2D adult best-match assignment were 81-99% in the 1-layer, 2-layer and 3-layer models with 1,000 nodes for astrocytes (Extended Data Figures 6D-F). This indicated that the MLP models had learnt to conduct highly accurate cell type assignment for some of the cell types in our training data with these parameters.

### Evaluation of the MLP models shows higher cross-2D/3D platform validation rates compared to label transfer cell type assignment

To evaluate the validation rates across the 2D/3D platforms for conducting cross-platform data integration, we used the MLP models that were trained on the 2D data to conduct cell type assignment on the 3D data (fetal and adult references). These MLP-assigned cell types were compared to the best-match assignment on the 3D data and the validation rate was defined as the number of cells with the same cell type assignment out of the total number of cells in the 3D data. We found that the maximum validation rate with the fetal reference was 25.1% and the maximum validation rate with the adult reference was 17.74%, as shown in Extended Data Figure 7.

To evaluate the per-cell type validation rates across platforms, we used the MLP models that were trained on the 2D data to conduct cell type assignment on the 3D data and calculated the percentages of MLP-assigned cells that agreed with the best-match assignment from the 3D data. We found that by mapping the fetal reference mapped data across 2D/3D platforms, the per-cell type validation rates were high at 44.1% to 58.7% for neuroepithelial cells (1-layer, 2-layer and 3-layer with 1,000 nodes), as shown in Extended Data Figure 8. Similarly, mapping the adult reference across 2D/3D platforms had high validation rates for some cell types, such as 61-74% for astrocytes (1-layer, 2-layer and 3-layer with 1,000 nodes). These validation rates were higher than the rates calculated by conducting Seurat label transfer cell type assignment across the 2D/3D platforms with the fetal reference (5.02% for neuroepithelial cells, range across all cell types = 3.13% to 50.62%) or adult reference (8.59% for astrocytes, range across all cell types = 0.36% to 18.21%), as shown in Extended Data Figure 4).

The validation rates with the 1-layer and 1,000-node MLP model ranged from 2.8% to 58.7% by mapping to the fetal reference (Extended Data Figure 8), which were similar to the rates calculated by conducting Seurat label transfer cell type assignment across the 2D/3D platforms with the fetal reference (ranging from 3.13% to 50.62%, Extended Data Figure 4). The validation rates with the 1-layer and 1,000-node MLP model ranged from 0.1% to 74.2% by mapping to the adult reference (Extended Data Figure 8), that were generally higher than the rates calculated by conducting Seurat label transfer cell type assignment with the adult reference (ranging from 0.36% to 18.21%, Extended Data Figure 4). These results indicate that the MLP models had higher accuracies by conducting cross 2D/3D cell type assignment with the adult reference, that is more relevant for studying neurodegenerative diseases such as AD.

### MLP models trained on 2D adult cell types had validation rates of 63.5% and 76.9% to a childhood brain dataset and a second adult brain dataset

We further evaluated the validation rates of applying these MLP models that were trained on our 2D dcOrg data with the best-match adult assignment (fetal/adult) to additional human brain tissue single-cell datasets. We assigned cell types in two new datasets – a childhood brain dataset^30^ and a second adult brain dataset^2^, and calculated the validation rates as the percentage of cells that were assigned by the MLP models to equivalent cell types as the reported cell types in the publications^2,30^. The highest validation rates by the MLP models trained on the 2D fetal data to assign cell types were 53.5% and 28% in the childhood and second adult data respectively, and the highest validation rates by the MLP models trained on the 2D adult data to assign cell types were 63.5% and 76.9% in the childhood and second adult data respectively (Extended Data Figure 9). These results indicate that there were high validation rates in data integration of dcOrgs and human adult brain tissues, by using these MLP models that were trained on 2D dcOrg data mapped to the adult reference for predicting cell types in human brain tissue scRNA-seq data from children or adults.

At the per cell type level, we found that the MLP models that were trained on the 2D fetal cell types had validation rates of 68.3% to 99.6% for astrocytes and neurons in the childhood dataset and validation rates of 40.6% to 66.8% for astrocytes and excitatory neurons in the second adult dataset (Extended Data Figures 10A-B). However, the validation rates for the other cell types were relatively lower in the childhood or second adult dataset. The MLP models that were trained on the 2D adult cell types had validation rates of 45.9% to 99.9% for astrocytes and neurons in the childhood dataset and validation rates of 19.6% to 99.6% for astrocytes and excitatory neurons in the second adult dataset (Extended Data Figures 10C-D). However, the validation rates for the other cell types were much higher in the childhood or second adult dataset when the MLP models were trained on the adult cells, compared to the validation rates for these cell types by the MLP models trained on the fetal cells. These results support our findings that MLP models that were trained on 2D dcOrg cells that were assigned to an adult reference, had higher validation rates across additional scRNA-seq data from human brain samples.

### Patterns of human and viral transcripts across the 2D dcOrgs and 3D cOrgs

We used the best-match assigned cell types for both the 2D dcOrgs and 3D cOrgs data for downstream analyses. We observed that some of the 3D organoid sections had more cells that were assigned to specific cell types, compared to other organoid sections (Extended Data Figures 11-12). There were cells that were assigned to all fetal or adult cell types across the 36 individual organoids that were used to generate the 3D cOrgs data. There were also similar sets of cells that were assigned to the same cell types in the fetal and adult references from the UMAP plots of the 2D and 3D data (Figures 2A-D), indicating that there are consistencies in cell type assignment to the fetal and adult references. We observed from the UMAP plots that were conducted with HSV-1 viral transcripts that all cell types were represented in the viral clusters in the 2D dcOrgs and 3D cOrgs (Figures 2E-H), indicating that the HSV-1 virus was able to infect all cell types found in the cerebral organoids in both the 2D and 3D systems (Extended Data Figures 13-16).

**Figure 2:**
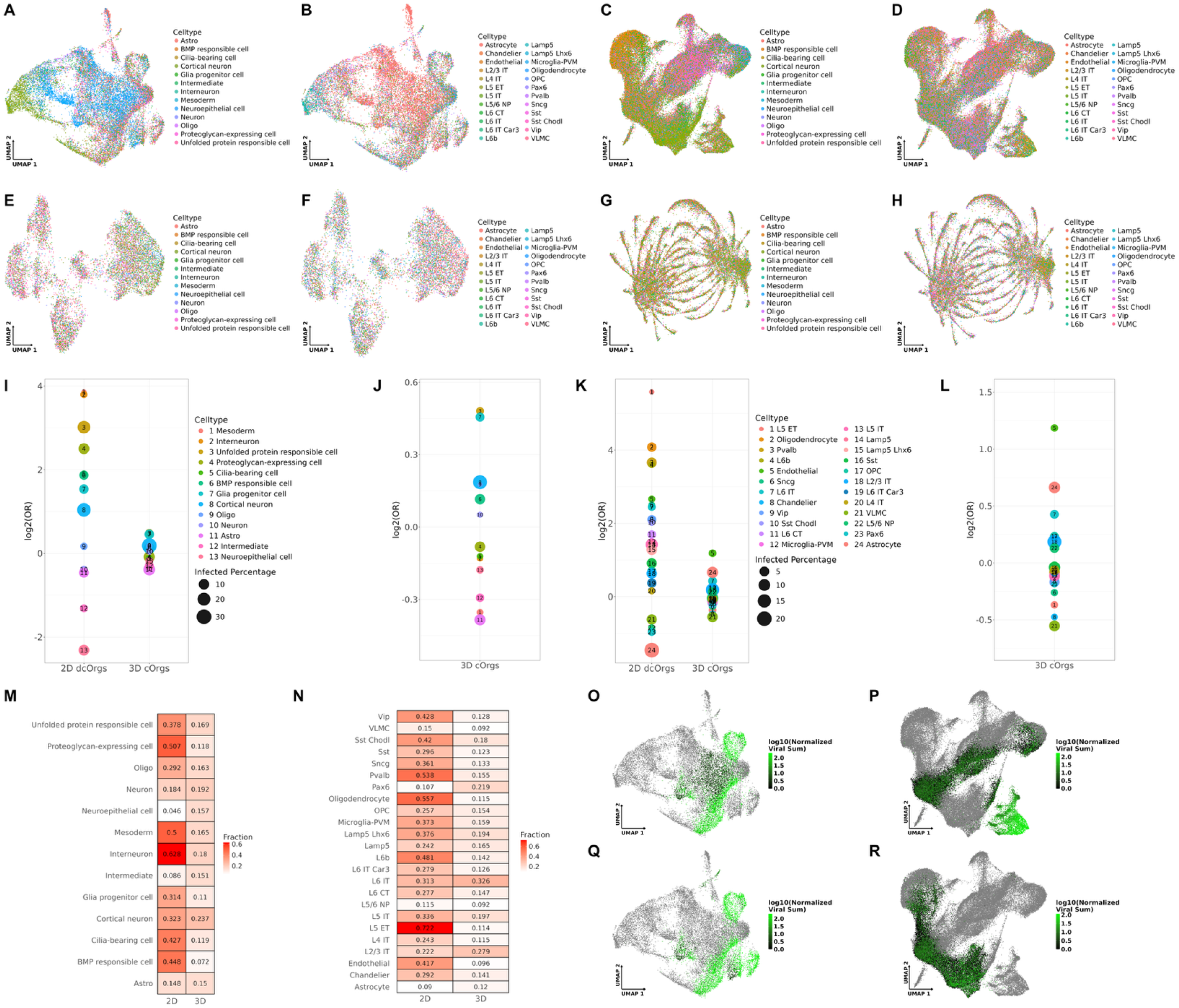
Distribution of human and viral transcripts, and cell type proportions across 2D dcOrgs and 3D cOrgs. (A) UMAP created by using the human transcripts in 2D scRNA-seq data that were mapped to fetal cell types. (B) UMAP created by using the human transcripts in 2D scRNA-seq data that were mapped to adult cell types. (C) UMAP created by using the human transcripts in 3D scRNA-seq data that were mapped to fetal cell types. (D) UMAP created by using the human transcripts in 3D scRNA-seq data that were mapped to adult cell types. (E) UMAP created by using the viral transcripts in 2D scRNA-seq data that were mapped to fetal cell types. (F) UMAP created by using the viral transcripts in 2D scRNA-seq data that were mapped to adult cell types. (G) UMAP created by using the viral transcripts in 3D scRNA-seq data that were mapped to fetal cell types. (H) UMAP created by using the viral transcripts in 3D scRNA-seq data that were mapped to adult cell types. (I) Fetal cell type proportion differences in 2D dcOrgs and 3D cOrgs, represented as *log_2_*(OR) and the sizes of the bubbles indicate the percentage of infected cells. (J) Magnified OR graph of the fetal cell type proportion differences in 3D cOrgs. (K) Adult cell type proportion differences in 2D dcOrgs and 3D cOrgs, represented as *log_2_*(OR) and the sizes of the bubbles indicate the percentage of infected cells. (L) Magnified OR graph of the adult cell type proportion differences in 3D cOrgs. (M) Heatmap depicting fractions of fetal cells with high normalized viral counts for each cell type in 2D dcOrgs and 3D cOrgs. (N) Heatmap depicting fractions of adult cells with high normalized viral counts for each cell type in 2D dcOrgs and 3D cOrgs. (O) UMAP with L2/3 IT neurons color-coded by normalized viral transcript counts from 2D dcOrgs. (P) UMAP with L2/3 IT neurons color-coded by normalized viral transcript counts from 3D cOrgs. (Q) UMAP with PVALB neurons color-coded by normalized viral transcript counts from 2D dcOrgs. (R) UMAP with PVALB neurons color-coded by normalized viral transcript counts from 3D cOrgs.

To evaluate the effects of HSV-1 infection on cell type proportion changes in the 2D dcOrgs and 3D cOrgs, we calculated the odds ratios (ORs) for each cell type, by computing the fraction of cell types in the HSV-1-infected samples over the fraction of cell types in the mock samples for the 2D and 3D data^8^. An OR>1 indicated that the cell type was enriched in the HSV-1-infected samples when compared to the mock samples and an OR<1 indicated that the cell type was depleted in the HSV-1-infected samples when compared to the mock samples. We found that there were more pronounced differences in cell type ORs in the 2D dcOrgs, compared to the cell type ORs in the 3D cOrgs (2D fetal ORs = [0.104, 14.662], 2D adult ORs = [0.196, 48.356], 3D fetal ORs = [0.73, 1.379], 3D adult ORs = [0.668, 2.273], Extended Data Figure 17).

To evaluate similarities and differences in cell type tropism for HSV-1 infection in 2D dcOrgs and 3D cOrgs, we compared the fractions of cells with high normalized viral counts for each cell type. We found that the cell types with high normalized viral counts within 2D dcOrgs were different from the cell types with high normalized viral counts in the 3D cOrgs (Figures 2M-R, Extended Data Figures 13-16), potentially indicating that the differences in the structure of the 3D cOrgs versus the 2D dcOrgs may lead to differences in viral replication.

### Temporal transcriptional expression of viral transcripts can be observed from the 2D and 3D scRNA-seq data

To further visualize the temporal transcriptomic expression of cell type clusters by viral transcript expression, we used a prior annotation of the HSV-1 viral transcripts that were categorized into 3 groups – group 1: intermediate early (IE) and early (E) transcripts, group 2: leaky late (LL) transcripts and group 3: true late (TL) transcripts^31^. For each viral cluster, we calculated the ORs for each group of transcripts to identify the enrichment or depletion of these 3 groups of viral transcripts within each viral cluster, with respect to the other viral clusters, and color-coded the viral clusters in the red, green and blue (RGB) channels respectively. We found that we were able to observe viral transcriptional dynamics in the 2D and 3D data (Figures 3A-B, Extended Data Figure 18). We color-coded the cell clusters in the viral UMAP plots by normalized viral counts and pseudotimes from the 2D dcOrgs and 3D cOrgs, and similarly, we found that the data could resolve temporal transcriptional dynamics and that trajectories from low to high viral counts aligned with trajectories from low to high pseudotimes in the 2D dcOrgs and 3D cOrgs data (Figures 3C-F). These results indicate that single-cell RNA sequencing (non-spatial and spatial) can be used to visualize and study viral transcriptional dynamics in human cerebral organoids (2D dcOrgs and 3D cOrgs).

**Figure 3:**
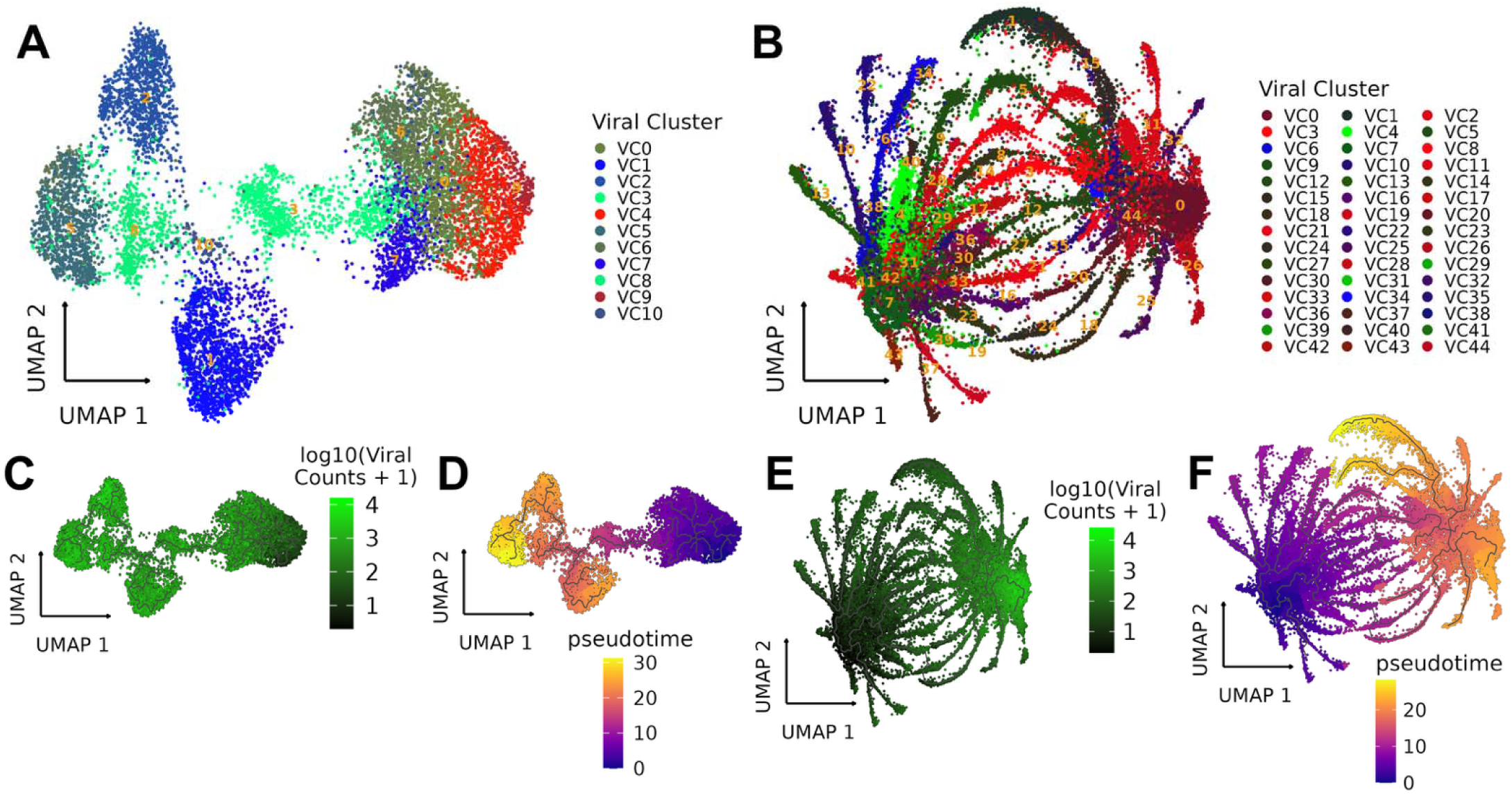
Pseudotime analyses on 2D dcOrgs and 3D cOrgs. (A) UMAP created by using the viral transcripts of infected cells in 2D dcOrgs and each cluster is color-coded by the enrichment of IE and E viral transcripts (red channel), LL viral transcripts (green channel) or TL viral transcripts (blue channel). (B) UMAP created by using the viral transcripts of infected cells in 3D cOrgs by using the viral transcripts and each cluster is color-coded by the enrichment of IE and E viral transcripts (red channel), LL viral transcripts (green channel) or TL viral transcripts (blue channel). (C) Pseudotime trajectories in 2D dcOrgs color-coded by the normalized viral transcript counts. (D) Pseudotime trajectories of infected cells in 2D dcOrgs rooted using cell clusters with the lowest viral transcript abundance. (E) Pseudotime trajectories in 3D cOrgs color-coded by the normalized viral transcript counts. (F) Pseudotime trajectories of infected cells in 3D cOrgs rooted using cell clusters with the lowest viral transcript abundance.

### Identifying cell types that were significantly clustered or dispersed revealed that endothelial cells and astrocytes were most significantly clustered in HSV-1-infected cOrgs, compared to mock cOrgs

To further evaluate if there may be cell types that were significantly clustered or dispersed due to HSV-1 infection in 3D cOrgs, we calculated the nearest neighbor (NN) distances for all cells in each cell type from the spatial data and evaluated the differences in the NN distances in the mock samples versus the infected samples by using Wilcoxon ranked sum tests (Figure 4A, Extended Data Figure 19). We observed that endothelial cells and astrocytes were the most significantly clustered cell types in the HSV-1-infected 3D cOrgs, compared to the mock 3D cOrgs (FDR=1.4×10^-40^ and 1.8×10^-37^ respectively). As an example, visualization of the physical locations of the astrocytes and endothelial cells assigned by the consensus approach is shown in Extended Data Figures 20-21. For comparison, visualization of the physical locations of microglia that did not have significantly different NN distances in the mock samples versus the infected samples (FDR=0.19), is shown in Extended Data Figure 22.

**Figure 4:**
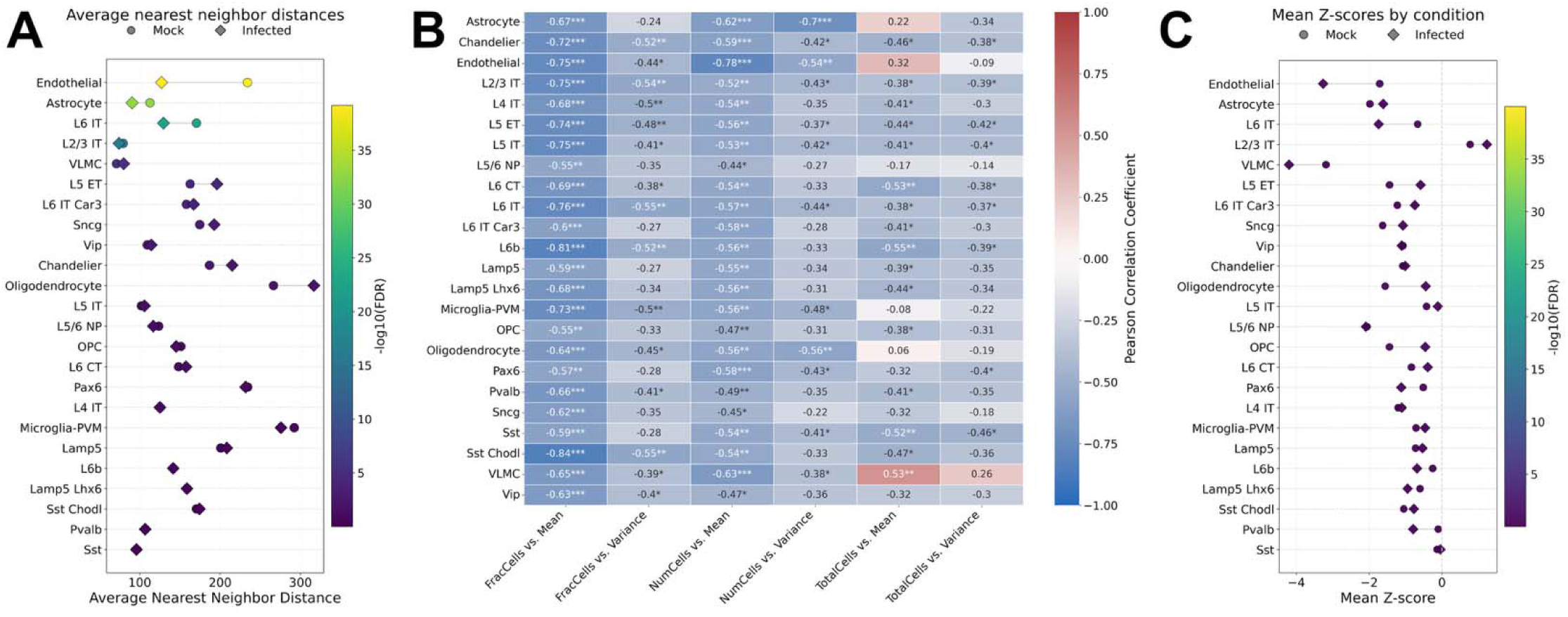
Observed and permuted cell-cell ANN distances for each cell type in infected cOrgs and mock cOrgs. (A) Plot of the observed ANN distances (*x*-axis) in infected cOrgs (represented by diamonds) and mock cOrgs (represented by circles) and color-coded by-log10(FDR) values, for each cell type (*y*-axis). (B) For each cell type (*y*-axis), the Pearson’s correlation coefficients between the fraction of cells in individual cOrgs, number of cells for a given cell type in individual cOrgs or total number of cells in individual cOrgs, and the mean ANN distance or variance in ANN distances from the 2,000 permutations (x-axis). (C) Plot of the mean observed Z-scores (mock or infected) calculated for the observed ANN distance with the permuted ANN distance distribution (*x*-axis) for each cell type (*y*-axis).

### Evaluating cell type specific spatial differences conditional on cell type proportion differences revealed clustering due to structural differences between HSV-1-infected cOrgs, compared to mock cOrgs

It is possible that the observed spatial differences observed between HSV-1-infected cOrgs and mock cOrgs were due to structural differences between the infected versus mock samples. For example, HSV-1 infection may have led to differences in the proportions of cell types within individual cOrg sections. If there were proportionally more endothelial cells in the HSV-1-infected cOrgs compared to the mock cOrgs, we may expect to find that the endothelial cells were more clustered in the HSV-1-infected cOrgs than the endothelial cells in the mock cOrgs. To evaluate this, we conducted a permutation test with 2,000 permutations to maintain the same density of cells for a given cell type in individual cOrgs. Next, we calculated the Pearson’s correlations within individual cOrgs between the total number of cells in the cOrgs, number of cells in each cell type in the cOrgs, fraction of cells for each cell type (as a measurement of density) in the cOrgs, and the average nearest neighbor (ANN), and the variances in NN distances (Figure 4B). We found that there were high anti-correlations between the fractions of cells and ANN distances, fractions of cells and variance in NN distances, number of cells and ANN distances, number of cells and variance in NN distances. Conversely, the total number of cells in the cOrgs was not highly anti-correlated. These results indicate that differences in the NN spatial distances observed in our samples may be driven by differences in the density of cells in HSV-1-infected cOrgs versus mock cOrgs.

To account for potential spatial localization differences due to overall differences in cell type density or fraction of cells in the infected cOrgs versus mock cOrgs, we calculated the Z-scores of the observed ANN compared to the permuted ANN distribution from the 2,000 permutations for each cOrg. We evaluated the differences in Z-scores between the infected cOrgs and mock cOrgs and observed that there were no significant FDR values for all cell types (Figure 4C, Extended Data Figure 24). These results indicate that there were no major structural differences in the mock cOrgs compared to infected cOrgs that may have led to potential overall differences in the observed NN distances. These analyses can be translated to human brain samples, for instance, to evaluate if there may be overall structural differences in the orientation of tissues, or selection of tissue samples, that may have led to NN spatial differences observed between brain samples from patients versus controls.

## DISCUSSION

The transcriptomics data by conducting scRNA-seq on cells within 2D dcOrgs and 3D cOrgs are different from the transcriptomics data by conducting scRNA-seq on cells from human fetal, childhood or adult brain samples. As such, there may be challenges in the use of spatial or non-spatial scRNA-seq data from cerebral organoid cells to infer cell types found in the human adult brain to study neurodegenerative diseases. We found that the use of MLP models were able to achieve similar validation rates as a current label transfer tool (Seurat) for cell type assignment to a fetal reference; however, the MLP models were able to achieve higher validation rates for cell type assignment to adult references, by using cross 2D/3D platform analyses to benchmark validation rates. These results demonstrate that the use of artificial neural networks and other sophisticated machine learning approaches to conduct cell type assignment on scRNA-seq data from complex human microphysiological systems to predict cell types found in the human adult brain may be a promising approach to studying neurodegenerative diseases.

Furthermore, we found that there were differences in HSV-1 infection in the 2D dcOrgs compared to HSV-1 infection in the 3D cOrgs, such as the magnitude in the proportions of cell types that were changed due to viral infections and cell type tropism for viral replication in the 2D versus 3D system. We found that the use of spatial and non-spatial scRNA-seq data from virus-infected 2D dcOrgs or 3D cOrgs could also enable the identification of specific viral clusters that could be used to track the pseudotime dynamics of the transcriptional life cycle of HSV-1 viral transcripts. Finally, we evaluated the nearest neighbor distances for pairs of cells for each cell type and found that cell types such as endothelial cells and astrocytes were significantly clustered in infected cOrgs versus mock cOrgs.

These methods, approaches and tools developed through our work can be applied broadly to data generated by other single-cell technologies (spatial or non-spatial). It will be interesting to use a sophisticated machine learning approach to further improve cell type assignment and to dissect cell cluster specific human-viral transcript interactions, given the sparse numbers of reads and transcripts detected using these unbiased, transcriptome-wide RNA-seq technologies. Furthermore, our methods can be further developed to integrate data from human *in-vitro* microphysiological systems, human brain samples, as well as samples from *in vivo* models, to gain insights into neurodegenerative diseases.

## MATERIALS AND METHODS

### Standard Protocol Approval

Research performed on samples and data of human and viral origin was conducted according to protocols approved by the Institutional Review Board (IRB) and Institutional Biosafety Committee (IBC) of UMass Chan Medical School. HSV-1 is a Biosafety Level (BSL) 2 pathogen.

### Source of human iPSCs and HSV-1 virus

The donor iPSC was the PGP1 participant from the Harvard Personal Genome Project (PGP) that had been characterized^8,22,32–36^. We used the HSV-1 K26GFP KOS strain^37,38^ that was provided by David Knipe and Hyung Suk Oh and we conducted infections similar to our prior protocols^8,39^, with some modifications.

### HSV-1 infection in 2D dcOrgs and 3D cOrgs

We differentiated 3D cerebral organoids using our methods based on protocols developed by Lancaster MA *et al*^8,21,40^. We used 5-month 2D dcOrgs for HSV-1 infection (Multiplicity of Infection, MOI of 2) as we had described previously^8^. For infections in 3D cOrgs, we first placed each individual 3-month intact cOrg in a single 1.5mL tube and washed the cOrgs twice in sterile ice-cold 1×DPBS, followed by a 10-minute spin-down each time. We assumed that there were 15,000 cells on the surface of the organoids and used an MOI of 10, with a 1-hour inoculation in a low volume 50μL of inoculation media with HSV-1, similar to previously reported protocols^41–43^. The cOrgs were then washed once with 1×DPBS and incubated in 2mL of differentiation media for viral replication to occur. Subsequently, after 47 hours, the cOrgs were then washed once with DPBS and fixed in 4% paraformaldehyde (PFA; Fisher Scientific 50-980-487) for 20 minutes, followed by another DPBS wash. The cOrgs were then passed through a sucrose gradient (10%, 20% and 30%) across another 2 days and 6 individual cOrgs were embedded in 1cm×1cm cryoblocks using Optimal Cutting Temperature (OCT) on dry ice. In total, we prepared 36 individual cOrgs (18 infected and 18 mock) and embedded 6 cOrgs of the same condition into a single cryoblock. One section was obtained from each cryoblock (10μm), leading to the generation of 6 independent sections for Stereo-seq sample preparation, library preparation and sequencing using DNBSEQ.

### Data processing

We used our 2D scRNA-seq data that we had previously generated and processed^8^. For our 3D scRNA-seq data, FASTQ files were processed through the SAW pipeline v6.0.2^44^ and we used custom R scripts for pre-processing of the data to integrate the human and viral transcript counts. The SAW workflow included mask splitting, barcode parameter generation, read mapping and generation of gene expression format (GEF) files. A split count of 16 and CID position 2_25 was used as recommended by the manufacturer, and a barcode length of 24bp, UMI length of 10 and one barcode mismatch allowed were used to generate barcode parameter files. Reads were mapped to a custom reference genome composed of GRCh38.p12 release 109, human alphaherpesvirus 1 Kos strain (GenBank: JQ673480.1) and a green fluorescent protein (EGFP) sequence with STAR alignment^45^. Gene counting was performed with the count module using the custom Gene Transfer Format (GTF) annotation file, enabling UMI correction (--umi_on, length = 10), with one barcode mismatch allowed and duplicate removal. Cell binning (with bin sizes of 50×50) to specify individual cells was conducted with the cellCut module to generate gene expression format (GEF) files. HSV-1 gene annotations were subsequently relabeled as duplicated viral gene names (e.g., RS1, RS1.1) to maintain consistency across datasets and duplicate HSV-1 features were removed prior to downstream analysis. Human and HSV-1 transcript count matrices were then integrated into combined objects by using Seurat v4.3.0^46^, with thresholds of min.cells = 5 and min.features = 100. We excluded cells with nFeature_RNA of <700 or>2,500 features. Counts were normalized with SCTransform with the glmGamPoi method and vst.flavor = “v2” in Seurat, or log-normalized for scPred and SingleR. Samples were integrated using Seurat’s SCT integration with 2,000 highly variable genes, similar to previously reported methods^47^.

### Cell type assignment

For the fetal reference, we used the data from a prior publication^23^. For the adult reference, we used the MTG dataset from the Allen Institute’s single-nucleus RNA-seq (snRNA-seq) atlas of the human middle temporal gyrus^24^. The data was initially down-sampled by using Seurat to 60,000 cells due to memory limitations, prior to integration and normalization. We conducted cell type assignment with Seurat v3 by using the TransferData function^25,26^, SingleR by using the de.method^27^, and scPred with a xgbDART based model and cell type assignment scores calculated using the “scPredict” function^28^. For the consensus approach, cell type assignment was determined by majority voting across the 3 tools. For the best-match approach, the cells that were unassigned by the consensus approach were assigned to the cell type assignment with the highest min-max scaled confidence score across the 3 tools.

### MLP model training by using leave-one-out approach

We used 9 biological replicates from the 2D scRNA-seq data for training the models by using 3 replicates of the mock dcOrgs (true uninfected or TU), and from the infected dcOrgs, we used 3 replicates of pseudo-uninfected (PU) cells and 3 replicates of true infected (TI) cells, as previously defined in our prior work^8^. The MLP models were trained by using 8 of the samples and the accuracies (number of correctly assigned cells divided by the total number of cells) were calculated from the remaining sample that was not used in the training. The following functions for the MLP models were used: rectified linear unit (ReLU) activation function for backpropagation, Adam optimizer with a fixed learning rate of 1×10⁻³, cross-entropy loss and full-batch mode for 50 epochs.

### Validation of MLP models by using additional datasets

For the childhood reference, we used a snRNA-seq dataset that was generated from cortical tissue samples obtained from children^30^. For the second adult reference, we used a snRNA-seq dataset that was generated from human post-mortem brain samples^2^. We mapped the cell types in the childhood reference to a set of cell types found in the fetal dataset and the cell types in the adult references, so that the cell types can be harmonized for validation (Extended Data Figure 25). Transcripts that were found in both the training and validation datasets were used. All cells in the fetal or adult references were used to train the MLP models in batches of 64 cells at a time, to reduce memory usage. In the second adult reference dataset, we used 500 cells from each cell type to calculate the validation rates, in order to avoid biasing the validation rates due to differences in the numbers of cells across cell types.

### Viral transcriptional life cycle categories

Intermediate early (IE) transcripts were UL54, US1, US12, US3 and RS1.1. Early transcripts were UL5, UL8, UL9, UL12, UL29, UL30, UL39, UL42 and UL52. Leaky late transcripts were UL18, UL19, UL35, UL26, UL32, UL21, UL36, UL41, UL46, UL48, UL49, US9, US10, UL27, US4, US6, US7, US8, UL45, UL11, UL34, RL1, US8A and UL24. True late transcripts were UL38, UL25, UL37, UL3, UL16, UL47, UL51, US2, US11, UL1, UL10, UL44, UL49A, US5, UL31, UL22 and UL2.

### Pseudotime trajectory analyses

Raw count matrices were converted into a cell_data_set (CDS) object using Monocle3 v1.3.1^48^. UMAP embeddings built previously from Seurat were transferred to the CDS object, and pseudotime was calculated by using graph learning and cells with the lowest abundance of viral transcripts were designated as the root state.

### Calculations of observed NN distances in mock cOrgs versus infected cOrgs

For each cell type, we calculated the NN distances for each cell that was assigned to the cell type in individual cOrgs. Rank sum tests were used to evaluate the NN distance distributions between mock cOrgs and infected cOrgs.

### Permutations to evaluate Z-scores of observed ANN distances

For each permutation, the coordinates of cells for a given cell type in each cOrg were randomly sampled from all coordinates for the cOrg. NN distances were calculated for the cells and the ANN distances were calculated across all permutations for individual cOrgs. The Z-scores of the observed ANN distances for individual cOrgs were calculated as 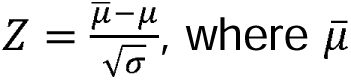 is the observed ANN distance in the cOrg, µ are the ANN distances calculated from the permutations and 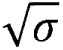 is the standard deviation of the ANN distances calculated from the permutations. We conducted t-tests on the Z-scores of the observed ANN distances between infected cOrgs and mock cOrgs, followed by FDR correction, to evaluate if there may be overall structural changes in the density of cell types between infected cOrgs and mock cOrgs.

### Schematic figures

All schematic figures in this manuscript were created using BioRender.

## ACKNOWLEDGEMENTS

This study is supported by startup funds (PIs: E.T.L. and Y.C.) and National Institutes of Health (NIH) grants R01AG083881 (PI: E.T.L.) and U01AG088673 (PI: Y.C.).

## AUTHOR CONTRIBUTIONS

E.T.L. planned and guided the project. A.D.D., P.D., Y.C. and E.T.L. conducted the research, analyses and methods development, with scripts generated by former lab members. P.D. and E.T.L. worked on quality control, consensus/best-match cell type assignment and downstream analyses. A.D.D. and E.T.L. worked on the MLP models. The HSV-1 viral stocks were provided by Hyung-Suk Oh and David M. Knipe. Cerebral organoid differentiation, infections and sample preparation were conducted by former lab members, guided by Y.C. and E.T.L. The list of viral transcripts in the transcription life cycle of HSV-1 were curated by Adrian R. Orszulak. E.T.L. wrote the initial manuscript with edits from all authors.

## DATA AVAILABILITY STATEMENT

Our scripts are uploaded to the project GitLab page (https://gitlab.com/elimlab/MipSScs). Our single-cell RNA-seq datasets will be uploaded to GEO.

